# The MKRN1-BRAF exon4-exon9 fusion is a targetable oncogenic driver

**DOI:** 10.1101/2021.07.22.453427

**Authors:** Megan A. Girtman, Craig S. Richmond, Paraic A. Kenny

## Abstract

**Background:** BRAF, when mutated at V600E, is a potent oncogenic driver in melanoma, lung and colorectal cancer with well understood signaling mechanisms and established treatment guidelines. Non-V600E mutations are less common, more functionally diverse and do not yet have clear treatment guidelines. One class of non-V600E mutations are BRAF fusion genes which typically involve the C-terminal kinase domain of BRAF joined to one of a wide repertoire of potential N-terminal fusion partners. Here, we functionally characterized an MKRN1-BRAF fusion gene which we detected in multi-gene panel sequencing of a metastatic colorectal tumor.

**Methods:** Levels of MEK/ERK pathway activity were evaluated by western blotting in HEK293 cells ectopically expressing MKRN1-BRAF or a series of other BRAF constructs. Dependence on dimerization was evaluated by introducing a dimerization deficiency mutation and drug sensitivity was evaluated by treatment with sorafenib, dabrafenib and trametinib.

**Results:** MKRN1-BRAF potently activated MEK/ERK signaling and did not require dimerization for activity. Among the inhibitors evaluated, trametinib most effectively suppressed MKRN1-BRAF driven pathway activity.

**Conclusion:** Our data demonstrate that the MKRN1-BRAF fusion gene encodes an oncoprotein that strongly activates MEK/ERK signaling in a trametinib-sensitive manner.

## INTRODUCTION

Metastatic colorectal cancer is the second and third most common cancer in women and men, respectively. There are 1.35 million new cases reported each year (Bray et al, 2018). Current therapies for the treatment of metastatic colorectal cancer include chemotherapy for eligible patients along with sequence analysis to determine the *KRAS, NRAS* and *BRAF* mutation status, which is primarily used to decide whether or not EGFR blocking antibodies (cetuximab or panitumumab) should be used in treatment (Biller & Schrag, 2021). Mismatch repair deficient tumors are treated with checkpoint immunotherapy (Le *et al*, 2020; Overman *et al*, 2017) and BRAF^V600E^ mutant tumors are treated with cetuximab and encorafenib (Tabernero *et al*, 2021).

Activating kinase gene fusions are infrequent in cancer but fusions involving NTRK (Cocco *et al*, 2018) and FGFR (Javle *et al*, 2018) genes exemplify the strong potential utility for targeted therapies for tumors in which these can be identified. Among colorectal cancer patients, kinase gene fusions are found in slightly less than 1% of cases but are enriched in tumors with high microsatellite instability (Cocco *et al*, 2019; Pagani *et al*, 2019). Targeted therapy for those patients, if they can be identified, might prove highly beneficial. Accordingly, it is important to study these mutations, determine which are true oncogenic drivers and, if so, which inhibitors might be clinically effective.

Recently, we evaluated an early 50s male patient having an *MKRN1-BRAF* fusion in a colorectal tumor that had metastasized to the liver. The fusion involved a breakpoint between *MKRN1* exon 4 and *BRAF* exon 9. This fusion protein retained the BRAF kinase domain but lacked the auto-inhibitory N-terminal region of BRAF. In this study, we ectopically expressed this MKRN1-BRAF fusion to determine the extent to which it may activate MEK/ERK signaling and evaluated a series of inhibitors for impact on this activation.

## MATERIAL AND METHODS

### Cell Culture

HEK293 cells were grown in Dulbecco’s Modified Eagle Medium (Thermo Fisher Scientific, Waltham, MA) supplemented with 10% fetal bovine serum (Gemini Bio-products, West Sacramento, CA). HEK293 cells were infected with lentivirus encoding MKRN1-BRAF, BRAF^WT^ and BRAF^V600E^. All BRAF cDNAs included a FLAG tag on the N-terminus.

### Antibodies

Antibodies against the following targets were used: phospho-p44/42 MAPK (ERK1/2) (#4370), p44/42 MAPK (ERK1/2) (#4695) phospho-MEK1/2 (#3958), Total MEK1/2 (#4694S) all purchased from Cell Signaling Technology (CST, Davers, MA), Raf-B (Santa Cruz Biotechnology, Dallas, TX) and FLAG M2 (Millipore Sigma, Burlington, MA).

### Inhibitors

Sorafenib, Dabrafenib and Trametinib were purchased from LC Laboratories (Woburn, MA). Drugs were freshly prepared by dissolving in DMSO (MilliporeSigma, Burlington, MA). Virally transduced cells were treated with 10 μM of drugs for 4 hours at 37°C.

### Western blot analysis

Cells were homogenized in Pierce RIPA lysis buffer (Thermo Fischer Scientific, Waltham, MA) containing protease and phosphatase inhibitors (1/100, Thermo Fischer Scientific, Waltham, MA). Lysates were quantified using a BCA assay (Thermo Fischer Scientific, Waltham, MA) and 35 μg of protein were loaded and size-fractionated by SDS-polyacrylamide gel electrophoresis (Bio-Rad Laboratories, Hercules, CA). Gels were then transferred to PVDF membranes. Membranes were blocked in 5% Blotto (Santa Cruz Biotechnology, Dallas, TX) for 60 minutes at room temperature and then incubated with the specific antibodies in dilution buffer at 4°C overnight. The blotted membranes were incubated with HRP-conjugated secondary antibodies (1:1000) at room temperature for 1hr. Protein expression level was detected and visualized using the Enhanced Chemiluminescence (ECL) detection system. (Thermo Fisher Scientific, Waltham, MA).

## RESULTS

In a genome sequencing panel report on an early 50s male patient with metastatic colorectal cancer, we identified a fusion gene consisting of exons 1-4 of *MKRN1* and 5-9 of *BRAF*. The fusion includes the BRAF kinase domain (Fig 1A), but lacks the N-terminal autoinhibitory domain of BRAF. While this pattern is generally consistent with prior observations on breakpoints in *BRAF* fusions with other partners (Ross *et al*, 2016), this particular fusion pair had not been functionally characterized.

**Fig 1.**
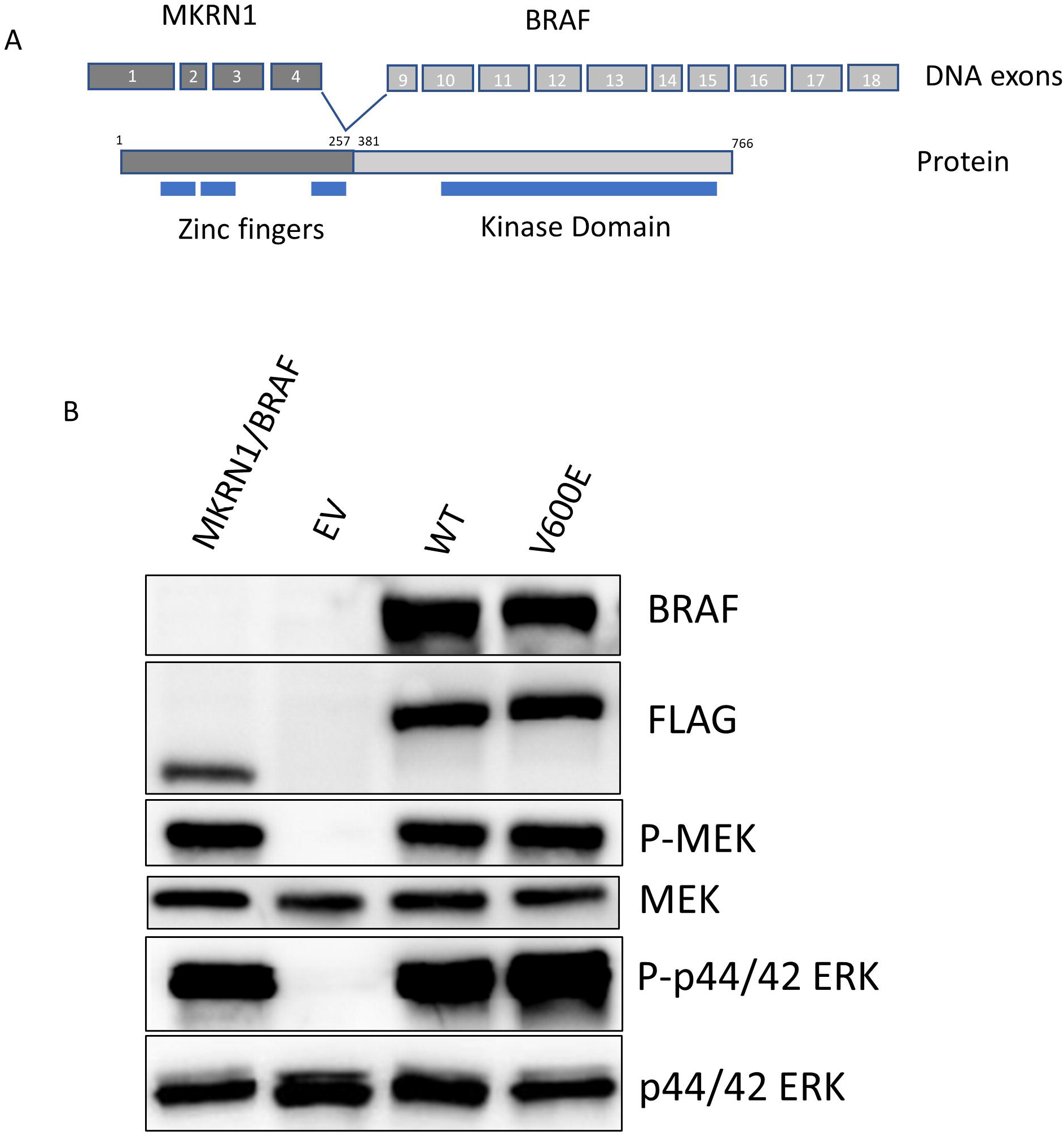
The MKRN1/BRAF fusion promotes MEK and ERK activity. A. Schematic representation of the fusion gene involving exons 1-4 of MKRN1 and 9-18 of BRAF, with key functional domains in the resulting fusion protein. B. Western blot analysis of pathway activity in HEK293 cells ectopically expressing the series of BRAF constructs.

Previous work has shown the value of ectopic expression in HEK293 and similar cell lines to characterize the function of BRAF fusion proteins (He et al, 2018; Jang et al, 2015; Tomic et al, 2017). We compared overall MEK/ERK pathway activity in cells expressing BRAF^WT^, MKRN1-BRAF or BRAF^V600E^, a class I mutant that elicits high MEK/ERK activity. Lysates were probed for BRAF, FLAG, pMEK, MEK, pERK, and ERK. In the fusion protein, the first 380 amino acids of BRAF are replaced by the first 257 amino acids of MKRN1, which eliminates the recognition epitope for the BRAF antibody (Fig 1B, top panel) and results in a size reduction of approximately 14 KDa, as seen from the anti-FLAG immunoblot (Fig 1B, second panel). Compared to empty vector transduced cells, cells expressing MKRN1-BRAF had increased MEK and ERK phosphorylation, similar to cells expressing BRAF^V600E^. These results demonstrate that this MKRN1-BRAF fusion activates MEK/ERK signaling at a level similar to the potent, BRAF^V600E^ oncogene.

One distinguishing feature that stratifies classes of BRAF mutants is a requirement for heterodimerization (Yao *et al*, 2017). Previous work has shown that the class I BRAF^V600E^ mutant activates the MEK/ERK pathway as a monomer. Class III mutants, such as BRAF^D594A^, require heterodimerization to function. To determine whether or not the MKRN1-BRAF fusion functions as a monomer or requires homo or heterodimerization in order to drive pathway activation, we generated dimerization deficient mutants by introducing a D509H mutation (Roring *et al*, 2012). Fig 2 shows levels of pathway activity in lysates from this series of HEK293 cell lines. Class I mutant BRAF^V600E^ shows similar levels of phosphorylated MEK and ERK with and without the D509H mutation, indicating the BRAF^V600E^ mutation is able to function as a monomer for downstream activation of MEK and ERK. In contrast, pathway activation by BRAF^WT^ overexpression was significantly attenuated by D509H. D509H introduction did not change the levels of pathway activation elicited by MKRN1-BRAF fusion, suggesting this fusion does not require dimerization mediated by its BRAF dimerization interface for signaling. While this indicates that it likely functions as a monomer, we cannot exclude the possibility that a dimerization event mediated by domains of the MKRN1 fusion component does not occur.

**Fig. 2.**
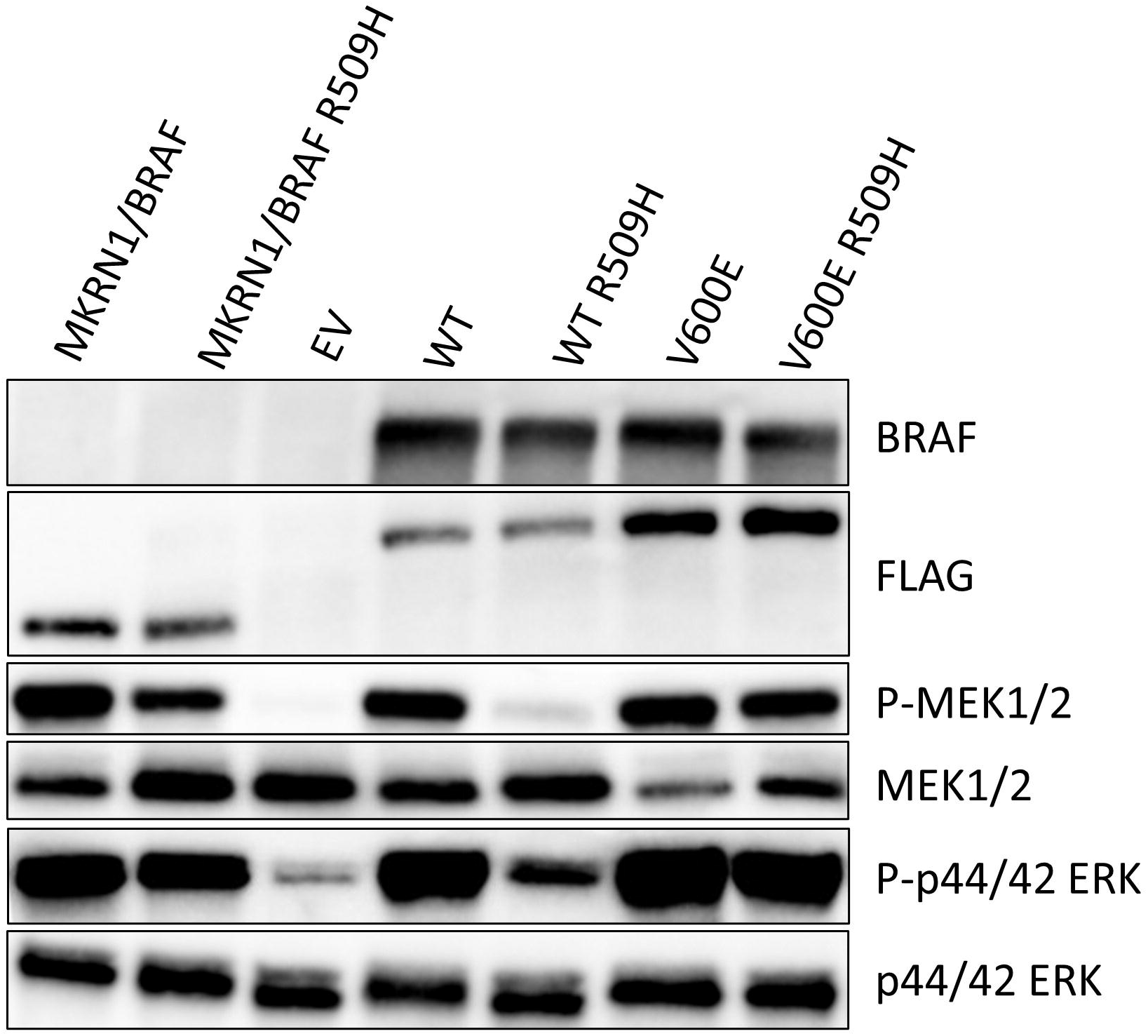
The MKRN1/BRAF fusion does not require the BRAF dimerization interface to promote MEK/ERK pathway activity. Western blot analysis of pathway activity in HEK293 cells ectopically expressing the series of BRAF constructs, with and without the D509H mutation which abrogates dimerization.

While therapeutic options for BRAF^V600E^ mutant cancers have been well established, the appropriate treatment strategy for non-classical BRAF mutations is still under investigation. We evaluated whether a pan-RAF inhibitor (sorafenib), mutant BRAF inhibitor (dabrafenib) and a MEK inhibitor (trametinib) could reduce the pathway activity levels elicited by the MKRN1-BRAF fusion in HEK293 cells (Fig 3A-C). The addition of sorafenib had only a very modest impact on MKRN1-BRAF driven MEK/ERK activity (Fig 3A). In contrast, dabrafenib had elicited a clear reduction of MEK/ERK activity, while the MEK inhibitor, Trametinib, reduced ERK activity to levels found in the empty vector transduced cells.

**Fig 3.**
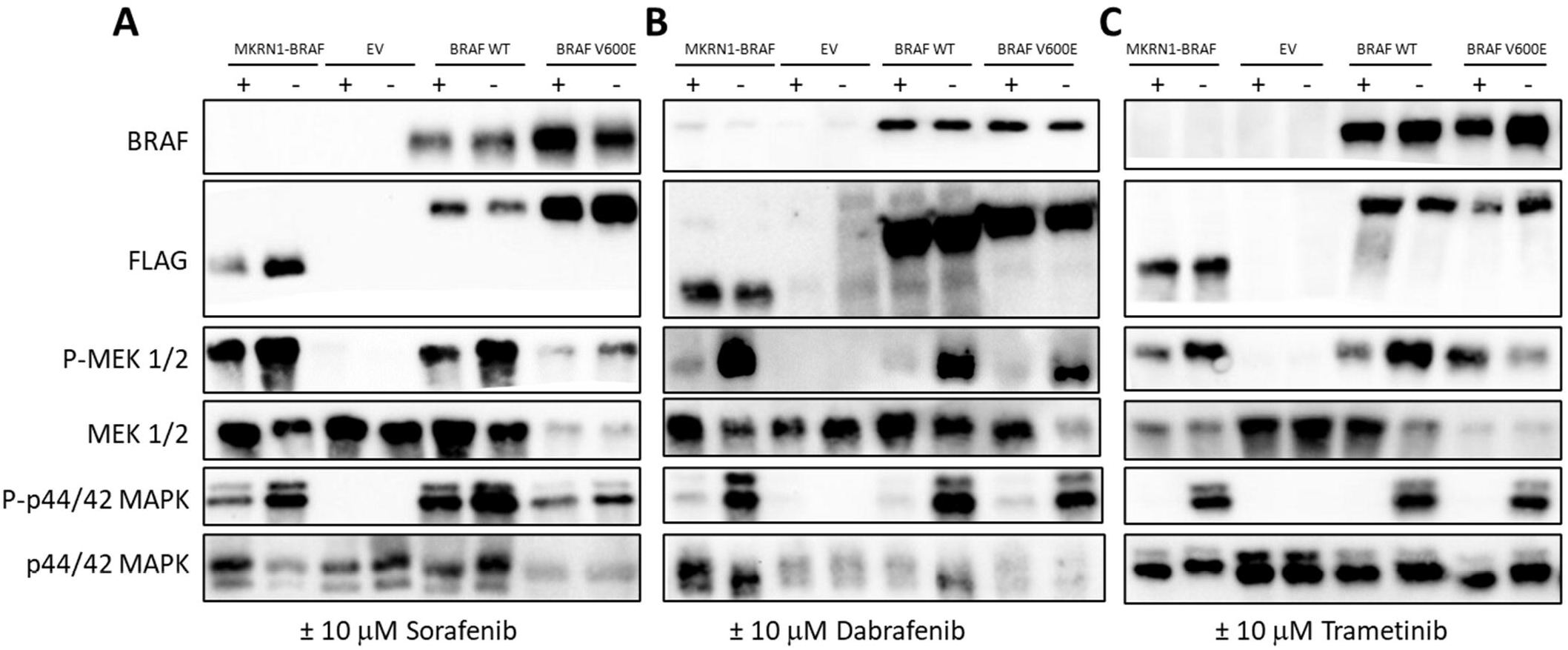
Among RAF/MEK/ERK pathway inhibitors, Trametinib most potently suppresses MEK/ERK pathway activation by MKRN1-BRAF. Western blot analysis of markers of MEK/ERK activation in HEK293 cells expressing the series of BRAF constructs after four hours of treatment with (A) Sorafenib, (B) Dabrafenib or (C) Trametinib.

## DISCUSSION

In this study, we functionally characterized the MKRN1-BRAF fusion gene, a recurrent although infrequent alteration in cancer. Ectopic expression of this fusion gene elicited MEK/ERK pathway activation at levels consistent with MKRN1-BRAF being an oncogene. Unlike BRAF^WT^, MKRN1-BRAF potently activated signaling as a monomer, in a manner similar to the well-characterized BRAF^V600E^ mutant. Evaluation of a series of FDA-approved inhibitors that modulate RAF/MEK/ERK signaling found that MEK inhibition with trametinib had the most potent attenuating effect on signaling induced by MKRN1-BRAF. Accordingly, we conclude that MKRN1-BRAF is an oncoprotein that is potentially therapeutically targetable.

Although trametinib inhibited MKRN1-BRAF driving signaling over a relatively short treatment time in culture (4 hours), clinical experience with trametinib in this setting has been mixed. Responses to trametinib in melanoma patients with PPFIBP2-BRAF (Menzies *et al*, 2015) and ZKSCAN1-BRAF (Ross *et al*., 2016) fusions have been reported. Subprotocol R of the NCI-MATCH trial evaluated trametinib in patients with non-V600E BRAF mutations. With 32 patients enrolled, this arm failed to meet its primary endpoint. Only a single BRAF fusion patient was enrolled in this arm and that patient experienced progressive disease (Johnson *et al*, 2020). Another Phase II study, restricted to advanced melanoma, enrolled two patients with BRAF fusions, one of whom experienced a partial response to trametinib (Nebhan *et al*, 2021). Trametinib had demonstrable clinical activity in pediatric low grade glioma patients with KIAA1549-BRAF fusions (Manoharan *et al*, 2020; Selt *et al*, 2020). Collectively, there appears to be a signal for efficacy in some settings but the small number of patients and tissue and fusion partner heterogeneity make it difficult to discern the potential impact of tumor type, fusion partner and co-occurring mutations on the likelihood of clinical benefit. In the patient in whom we detected the *MKRN1-BRAF* fusion, the co-occurring mutations (*EGFR, FGFR1* and *MYC* amplification, *TP53* G245S) limited clinical enthusiasm for attempting to target BRAF.

In general, the potential additional gain/loss-of-function consequences attributable to disruption of the fusion partner of BRAF is under explored. While removal of the N-terminal autoinhibitory domain of BRAF, thus allowing unrestrained kinase activity, is almost certainly the major oncogenic effect of the MKRN1-BRAF fusion, the potential impact of MKRN1 dose-reduction resulting from capture of one allele in the BRAF fusion also warrants discussion. MKRN1, also known as Makorin, is an E3 ubiquitin ligase. MKRN1 substrates include the EAG1 potassium channel (Fang *et al*, 2021), the catalytic subunit of telomerase, hTERT (Kim *et al*, 2005), as well as p53 and p21 (Lee *et al*, 2009). Experimental depletion of MKRN1 resulted in increased telomerase activity (Kim *et al*., 2005) and suppression of DNA damage induced cell death (Lee *et al*., 2009), either of which could confer additional advantages beyond activation of MEK/ERK signaling during cellular transformation or subsequent therapy.

Given the enrichment for BRAF fusion mutations among colorectal tumors with mismatch repair deficiency, many of these patients are likely to first receive checkpoint immunotherapy as standard-of-care given the well-established survival benefit (Le *et al*., 2020; Overman *et al*., 2017), leaving consideration of MEK inhibition or another strategy for a later line of treatment. IMblaze370 evaluated the combination of another checkpoint immunotherapy, atezolizumab, and a different MEK inhibitor, cobimetinib, in patients almost exclusively mismatch repair proficient and BRAF^WT^ colorectal cancers (Eng *et al*, 2019). Although the combination was not beneficial in that particular population, the MEK and checkpoint inhibitor regimen was generally well tolerated, raising the possibility of evaluating whether it might have higher activity in mismatch repair deficient / BRAF-fusion tumors.

## ACKNOWLEDGEMENTS

This work was supported by the Gundersen Medical Foundation. PAK holds the Dr. Jon & Betty Kabara Endowed Chair in Precision Oncology. MAG was supported by the Norman L. Gillette Jr Postdoctoral Research Fellowship.

